# Transcriptional adaptation of drug-tolerant *Mycobacterium tuberculosis* in mice

**DOI:** 10.1101/2023.03.06.531356

**Authors:** Elizabeth A Wynn, Christian Dide-Agossou, Matthew Reichlen, Karen Rossmassler, Reem Al Mubarak, Justin J Reid, Samuel T Tabor, Sarah E M Born, Monica R Ransom, Rebecca M Davidson, Kendra N Walton, Jeanne B Benoit, Amanda Hoppers, Allison A Bauman, Lisa M Massoudi, Gregory Dolganov, Payam Nahid, Martin I Voskuil, Gregory T Robertson, Camille M Moore, Nicholas D Walter

**Affiliations:** Rocky Mountain Regional VA Medical Center, Aurora, CO, USA; Department of Biostatistics and Informatics, University of Colorado, Anschutz Medical Campus, Aurora, CO, USA; Consortium for Applied Microbial Metrics, Aurora, CO, USA; Division of Pulmonary Sciences and Critical Care Medicine, University of Colorado Anschutz Medical Campus, Aurora, CO, USA; Department of Immunology and Microbiology, University of Colorado Anschutz Medical Campus, Aurora, CO, USA; Division of Hematology, Department of Medicine, University of Colorado School of Medicine, Aurora, CO, USA; Center for Genes, Environment and Health, National Jewish Health, Denver, CO, USA; Mycobacteria Research Laboratories, Department of Microbiology, Immunology, and Pathology, Colorado State University, Fort Collins, CO, USA; Division of Infectious Diseases and Geographic Medicine, Stanford University, Palo Alto, CA, USA; Division of Pulmonary and Critical Care Medicine, University of California San Francisco, CA, USA; UCSF Center for Tuberculosis, University of California, San Francisco, CA, USA

## Abstract

Transcriptome evaluation of *Mycobacterium tuberculosis* in the lungs of laboratory animals during long-term treatment has been limited by extremely low abundance of bacterial mRNA relative to eukaryotic RNA. Here we report a targeted amplification RNA sequencing method called SEARCH-TB. After confirming that SEARCH-TB recapitulates conventional RNA-seq *in vitro*, we applied SEARCH-TB to *Mycobacterium tuberculosis-*infected BALB/c mice treated for up to 28 days with the global standard isoniazid, rifampin, pyrazinamide, and ethambutol regimen. We compared results in mice with 8-day exposure to the same regimen *in vitro*. After treatment of mice for 28 days, SEARCH-TB suggested broad suppression of genes associated with bacterial growth, transcription, translation, synthesis of rRNA proteins and immunogenic secretory peptides. Adaptation of drug-stressed *Mycobacterium tuberculosis* appeared to include a metabolic transition from ATP-maximizing respiration towards lower-efficiency pathways, modification and recycling of cell wall components, large-scale regulatory reprogramming, and reconfiguration of efflux pumps expression. Despite markedly different expression at pre-treatment baseline, murine and *in vitro* samples had broadly similar transcriptional change during treatment. The differences observed likely indicate the importance of immunity and pharmacokinetics in the mouse. By elucidating the long-term effect of tuberculosis treatment on bacterial cellular processes *in vivo*, SEARCH-TB represents a highly granular pharmacodynamic monitoring tool with potential to enhance evaluation of new regimens and thereby accelerate progress towards a new generation of more effective tuberculosis treatment.

## INTRODUCTION

Tuberculosis (TB) is an ongoing public health crisis, killing approximately 1.2 million people each year.^1^ A key challenge for global TB control is the length of treatment required to cure TB, 4-6 months for infection with drug-sensitive genotypes. The need for prolonged treatment is thought to be due to a transition of the *M. tuberculosis* (*Mtb*) population to harder-to-kill phenotypes as therapy progresses.^2^ At the start of treatment, the *Mtb* population is highly susceptible to killing. When mice or humans initiate the global standard isoniazid, rifampin, pyrazinamide, ethambutol (HRZE) regimen, ∼99% of the culturable bacterial population is eliminated during the initial days to weeks of the bactericidal phase. Thereafter, the rate of killing slows considerably^3–7^ as drug-tolerant phenotypes come to dominate the residual *Mtb* population. Drug-tolerant phenotypes have been defined as *Mtb* with decreased susceptibility to killing despite an absence of drug resistance conferring mutations.^2, 8^

Eliminating drug-tolerant *Mtb* phenotypes is considered crucial to shortening the duration of TB treatment.^2^ The central focus of contemporary drug discovery and regimen development is drug-tolerant *Mtb* phenotypes that cause relapse and dictate the need for prolonged treatment.^2, 9–11^ Unfortunately, the physiologic state of the *Mtb* subpopulation that withstands drug treatment *in vivo* remains obscure, limiting our ability to rationally design regimens that target specific cellular processes.

Transcriptional profiling has been used extensively as a readout of how drugs affect bacterial cellular processes *in vitro.*^12–18^ By contrast, we are not aware of previous genome-wide transcriptional profiling of *Mtb* during prolonged treatment of laboratory animals*. In vivo* analyses are important because the pathogen adapts its cellular processes to the physicochemical conditions and immune responses it encounters in the host, resulting in a phenotype and transcriptome distinct from that measured *in vitro*.^19–27^ Since phenotypic characteristics of the *Mtb* population strongly influence drug sensitivity,^2^^8^ understanding the effect of drugs on the bacterial populations that cause disease *in vivo* would enable us to better “know the enemy,” thereby informing drug and regimen development.

The central impediment to transcriptional profiling *in vivo* is that *Mtb* mRNA is generally present at extremely low abundance relative to host RNA even in a treatment-naïve animal. When the pathogen burden decreases due to treatment, quantification of *Mtb* mRNA becomes progressively more difficult. To elucidate *Mtb* signal from the overwhelming eukaryotic background, diverse methods have been used to first enrich and then quantify *Mtb* mRNA (**Fig. 1**). Enrichment methods have included various combinations of differential cell lysis to separate *Mtb* and eukaryotic material during sample preparation, reverse transcription with pathogen-specific primers, depletion of host and/or bacterial rRNA, various hybridization capture methods, and selective PCR amplification of target *Mtb* transcripts. Enrichment is followed by quantification via microarray,^19, 20, 27, 29^ multiplex PCR,^7, 21, 25, 30–33^ or RNA-seq.^22, 26, 34–37^ Unfortunately, even the most sophisticated and contemporary platforms have limited sensitivity that has precluded transcriptional evaluation of the effect of long-term treatment with potent TB regimens. In this work, we demonstrate SEquening after Amplicon enRiCHment for TB (SEARCH-TB), a targeted, highly sensitive, pathogen-specific sequencing method for enriching and quantifying the *Mtb* transcriptome in tissues of infected animals. Using a novel combination of enrichment (differential cell lysis during sample preparation + targeted amplification) and quantification (RNA-seq), SEARCH-TB achieves sensitivity that will enable routine use of transcriptional profiling for drug evaluation in animal models, expanding understanding of drug effect and accelerating new regimen development.

**Fig. 1.**
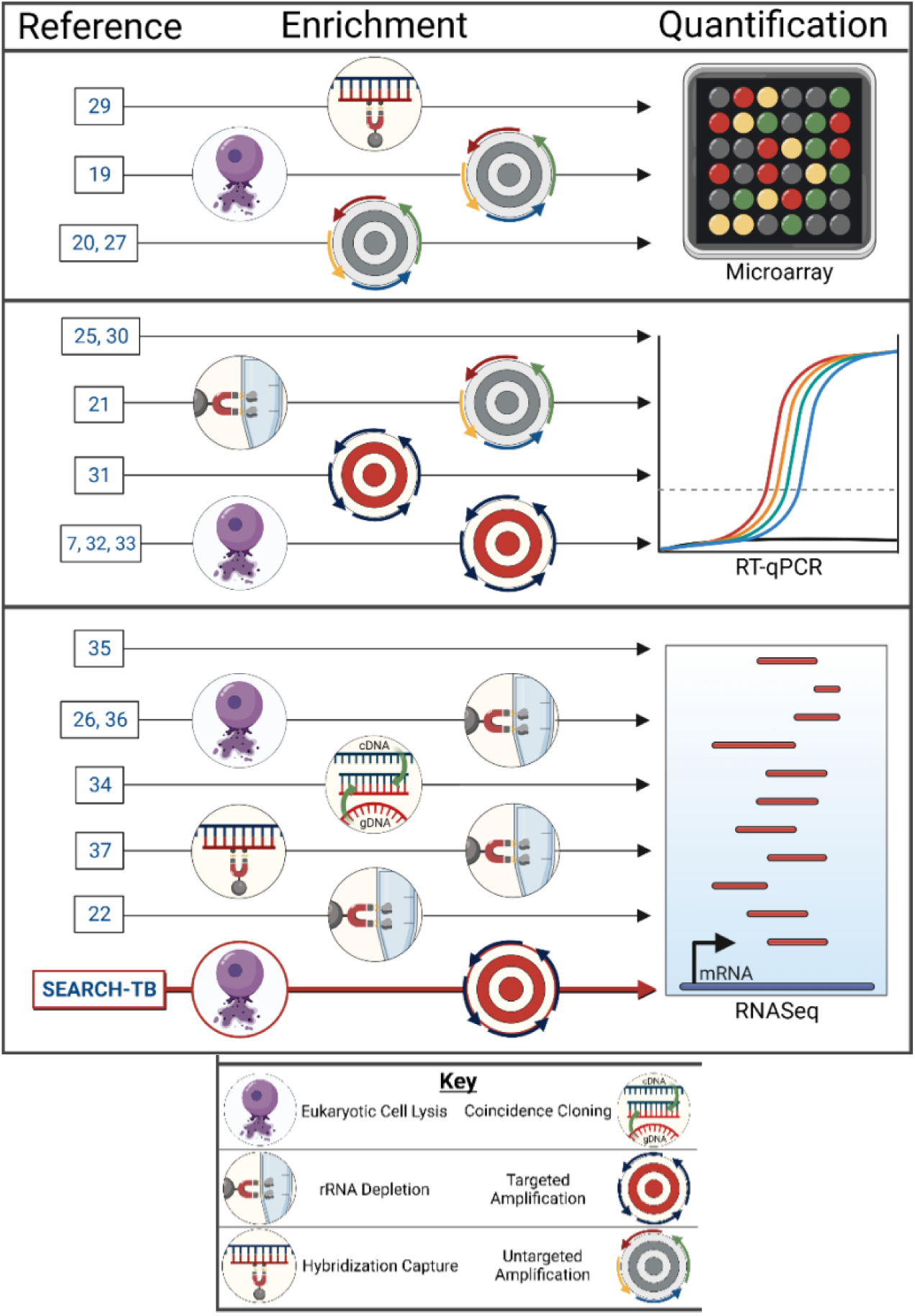
Visual summary of methods previously used to quantify the *Mtb* transcriptome *in vivo*. Each horizontal arrow represents a distinct combination of enrichment and quantification. Varying enrichment methods are represented via the symbols shown in the key. SEARCH-TB is a unique combination of enrichment (eukaryotic cell lysis + targeted amplification) followed by quantification via RNA-seq that has enabled transcriptome evaluation in mice treated for weeks with a potent combination regimen. Image created with Biorender.com.

Here, we used SEARCH-TB to evaluate gene expression of the *Mtb* population that persists in the lungs of BALB/c mice treated with HRZE for up to four weeks. We further used SEARCH-TB to compare the effect of HRZE *in vivo* and *in vitro.* SEARCH-TB has the potential to serve as a highly granular readout of drug effect by providing a molecular readout of cellular adaptations of *in vivo* drug-tolerant *Mtb*.

## METHODS

### Design of the SEARCH-TB assay

The SEARCH-TB assay uses an AmpliSeq for Illumina custom pool designed to amplify coding sequence (CDS) of *Mtb* complex (MTBC) organisms. To assure amplification across diverse lineages, the design used eight MTBC reference genomes (**Table S1**). Annotations were prepared for *Mtb* H37Rv (NC_000962) and *Mtb* Erdman (AP012340.1) as described in the Supplemental Information. To avoid off-target amplification (*i.e.*, of non-MTBC organisms), primers were cross-referenced during design with 12 “exclusion” genomes that included phylogenetically diverse bacteria as well as human and mouse (**Table S2**).

### Evaluation of amplification bias

We tested for amplification bias (*i.e.,* differences in amplification efficiency between primer pairs targeting different *Mtb* sequences) using replicate human lung RNA samples spiked with *Mtb* genomic DNA (gDNA) (Supplemental Information). Since all targeted sequences are present as single copies in gDNA, an entirely unbiased assay would hypothetically result in the same copy number for all targets, indicating that all primer pairs amplified with identical efficiency. We defined amplification bias as deviation from this ideal by comparing the observed expression for a gene to the expected expression assuming no amplification bias (Supplemental Information).

### Evaluation of repeatability of amplification

To evaluate repeatability, we spiked 1pg *Mtb* RNA into 1ng human lung RNA (Supplemental Information). We compared counts per million (CPM) values for each gene between technical replicates. We also evaluated repeatability over time by prepping and sequencing 19 of the *in vitro* and murine samples described below two times with up to a 4-month intervening interval. We quantified batch effect by calculating the difference in expression (normalized with DESeq2’s variance stabilizing transformation^38^) of each gene between replicate pairs, then averaging across replicates. We compared the magnitude of the batch effect with the observed treatment effect.

### Concordance of SEARCH-TB with conventional RNA-seq

We evaluated whether SEARCH-TB identified the same transcriptional changes as a conventional RNA-seq method without *Mtb* targeted amplification (Illumina TrueSeq) after 24-hour *in vitro* isoniazid (INH) exposure (Supplemental Information). We first evaluated RNA from control (N=4) and INH-treated samples (N=4) via conventional RNA-seq to serve as a reference standard. We then spiked the same RNA from control and INH-treated *Mtb* into human lung RNA at a ratio of 1:1,000 and sequenced via SEARCH-TB. After calculating differential expression between control and INH-treated samples separately for conventional RNA-seq and SEARCH-TB using edgeR,^39^ we compared the significant genes and fold-changes identified by the two platforms.

### *In vitro* experiments

*Mtb* strains H37Rv and Erdman were cultured *in vitro* using Middlebrook 7H9 broth (Difco) supplemented with 0.085 g/l NaCl, 0.2% glucose, 0.2% glycerol, 0.5% BSA, and 0.05% Tween-80. All culturing was performed at 36.5°C and 5.0% CO_2_. Single use frozen *Mtb* aliquots were revived in 7H9 and grown to mid-log phase then cultures were diluted to OD_600_=0.05, dispensed in 5.0 ml aliquots into sterile glass tubes (20 by 125 mm) containing sterile stir bars (12 by 4.5 mm), and outgrown for 18 h under rapid agitation (∼200 rpm stirring speed using a rotary magnetic tumble stirrer) prior to the initiation of drug exposure. RNA was collected from *Mtb* H37Rv exposed to INH or *Mtb* Erdman exposed to HRZE *in vitro* (Supplemental Information).

### Murine drug experiments

All animal procedures were conducted according to relevant national and international guidelines and approved by the Colorado State University Animal Care and Use Committee as described in the Supplemental Information. Briefly, female BALB/c mice, 6 to 8 weeks old, were aerosol infected (Glas-Col) with *Mtb* Erdman strain resulting in the deposition of 4.55±0.03 (SEM) log10 CFU in lungs one day following aerosol. After 11 days, five mice were euthanized to serve as the pre-treatment control group. Groups of five mice each were treated with HRZE at standard doses five days a week for 14 or 28 days before euthanasia. Lungs were aseptically dissected, and flash frozen in liquid nitrogen before processing.

### RNA extraction, sequencing, and data preparation

RNA extraction, library preparation, sequencing and data preparation are detailed in the Supplemental Information.

### Statistical analysis of murine and *in vitro* experiments

Murine and *in vitro* sequence data were analyzed together using edgeR^39, 40^ to identify the effect of HRZE treatment in and compare gene expression between murine and *in vitro* experiments. We fit negative binomial generalized linear models to each gene and included terms for murine and *in vitro* time points (control, day 14 and day 28 in mice; control, day 4 and day 8 *in vitro*). Likelihood ratio tests were performed to compare expression between murine time points, between *in vitro* time points, and between murine and *in vitro* experiments before and at the end of treatment. Genes with Benjamini-Hochberg adjusted *P*-value^41^ less than 0.05 were considered significant. For murine experiments, we used hierarchical clustering to identify groups of genes with similar changes in gene expression over the course of treatment as follows. First, for genes that were differentially expressed between at least two time points, we calculated the expected expression at each time point using the edgeR models. Then, the expected expression values were hierarchically clustered based on Euclidian distance using Ward’s method^42^ to find clusters of genes with similar patterns of expression over time.

Principal component analysis (PCA) was performed using data for the 500 most variable genes after normalization with DESeq2’s variance stabilizing transformation.^38^

Using hypergeometric tests in the hypeR R package,^43^ we performed functional enrichment for each pairwise combination of murine and *in vitro* time points to evaluate whether differentially expressed genes were overrepresented in gene categories established by Cole^44^ or curated from the literature (**Table S3**). Enrichment analysis was run twice for each pairwise combination, first using significantly upregulated genes and then using significantly downregulated genes. Gene categories with <8 genes were excluded. Gene categories with Benjamini-Hochberg adjusted *P*-values^41^ less than 0.05 were considered significant. All analysis used R (v4.1.1).^45^

### Online analysis tool

Differential expression, functional enrichment, and visualizations can be evaluated interactively using an Online Analysis Tool [https://microbialmetrics.org/analysis-tools/] created using the R package Shiny.^46^

## RESULTS

### Validation of SEARCH-TB assay

#### Results of SEARCH-TB design

Primers were designed to amplify 3,733 (92.6%) of 4,031 CDS in the *Mtb* H37Rv genome. Primers for 95 genes failed to amplify in initial testing and were excluded from analysis.

Additionally, we excluded 70 genes that were not present in the *Mtb* Erdman strain used in murine and *in vitro* experiments. The final panel analyzed targeted 3,568 *Mtb* genes (**Fig. S2**).

#### Evaluation of amplification bias

Using gDNA (a matrix in which each target sequence should be present in equal abundance), we quantified deviation from the ideal value that would be expected if all primers had identical amplification efficiency (**Fig. 2a**). Of all primer pairs, 78% were within one log_2_ fold change of the ideal value.

**Fig. 2.**
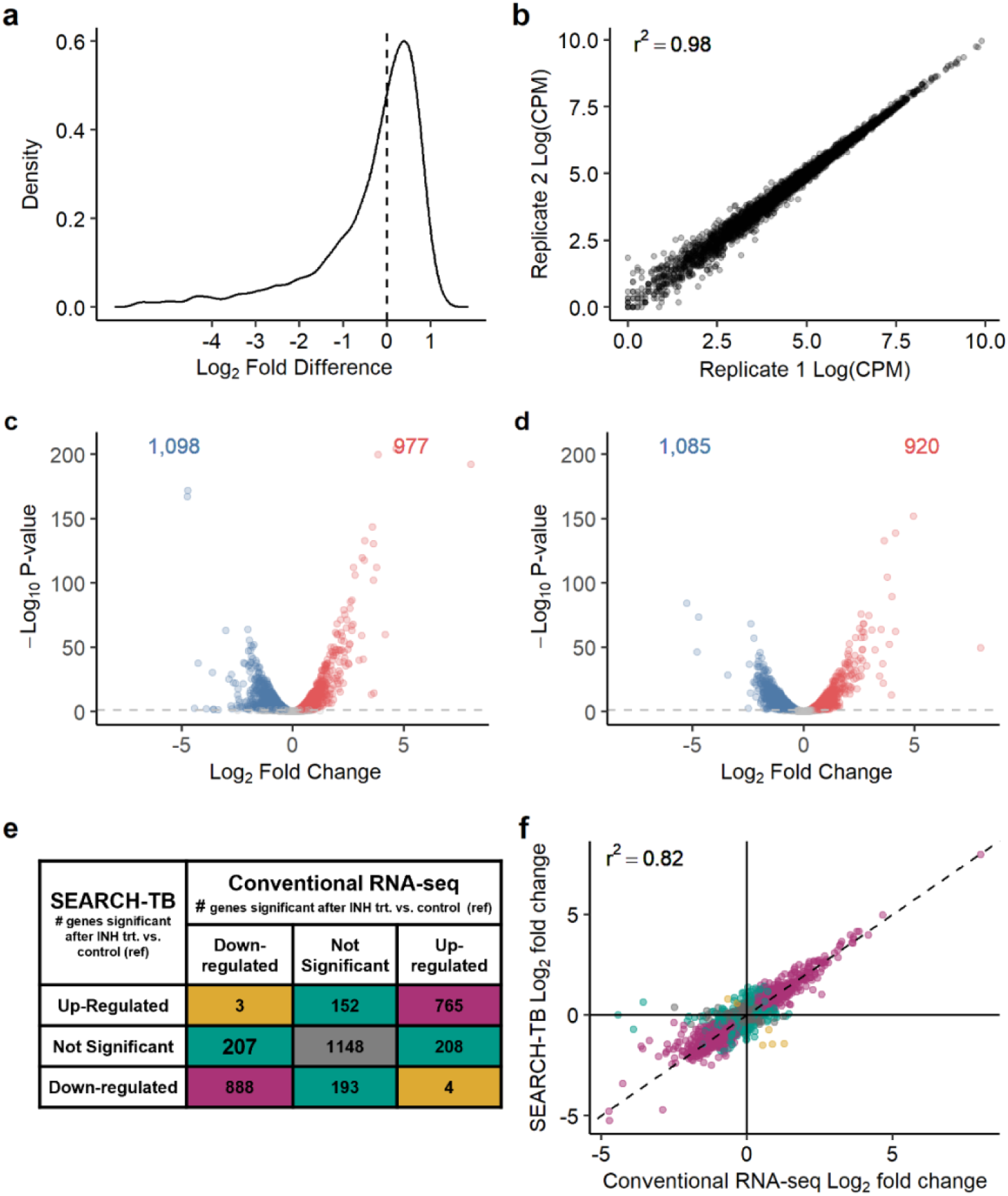
Evaluation of SEARCH-TB platform. a. Distribution of log2 fold differences of normalized gDNA SEARCH-TB expression data relative to the value expected if there were no amplification bias. Zero (dashed vertical line) represents no amplification bias. **b.** Evaluation of repeatability of SEARCH-TB showing normalized expression data (log counts per million) for two technical replicates in which *Mtb* RNA was spiked into human lung RNA. **c-d.** Volcano plot showing log2 fold-changes and -log10 *p-*values induced by *in vitro* INH exposure as quantified by RNA-seq (**c**) and SEARCH-TB (**d**). Genes significantly down- and upregulated with INH exposure relative to control (adj. p-value< 0.05) are shown in blue and red. **e.** Comparison of differential expression between INH treated samples and control samples from RNA-seq or SEARCH-TB data. Purple shading indicates genes with concordant fold-change direction and significance between RNA-seq and SEARCH-TB. green shading indicates genes that were significant in RNA-seq or SEARCH-TB results, but not both. Gold shading indicates genes that were significant for both RNA-seq and SEARCH-TB, but in opposite directions. Gray shading indicates genes that were not significantly differentially expressed in either RNA-seq or SEARCH-TB. **f.** Comparison of INH vs. control fold-changes from RNA-seq data vs. SEARCH-TB data. Purple, green, gold and gray colors have the same meaning as in **e**.

#### Repeatability of amplification

SEARCH-TB results were highly repeatable among spike-in technical replicates (**Fig. 2b**). The batch effect between replicates prepared and sequenced over time was minimal relative to the treatment effect (**Fig. S3**).

#### Concordance of SEARCH-TB with conventional RNA-seq

RNA from *in vitro* exposure showed a similar direction and scale of differential expression when sequenced via SEARCH-TB and conventional RNA-seq (**Fig. 2c-d**). The gene sets identified as differentially expressed by the two platforms strongly overlapped (**Fig. 2e**). The fold-changes quantified by the two platforms showed strong agreement (*R*^2^=0.82) (**Fig. 2f**), indicating that while SEARCH-TB is uniquely capable of profiling in drug-treated animals, both platforms provide the same biological information in *in vitro* RNA.

### Transcriptional differences of between *Mtb* in murine and *in vitro* controls

In the absence of HRZE, a PCA plot showed the transcriptional profiles of the pre-treatment murine control and the *in vitro* early log-phase growth control were distinct (**Fig. 3a**) with 2,444 (68%) differentially expressed genes (**Fig. 3b**). Compared with the *in vitro* control, untreated mice appeared to harbor a less active *Mtb* phenotype with lower expression of genes associated with rRNA protein synthesis (adj-*P*=1.7×10^-18^), aerobic metabolism (adj-*P*=2.1×10^-5^), fatty acid synthesis (adj-*P*=0.002) and other growth-associated processes (Online Analysis Tool). Conversely, *Mtb* in untreated mice had higher expression of processes previously associated with adaptation to the host^47^, including genes in the DosR regulon that responds to hypoxia, nitric oxide, and carbon monoxide^48, 49^ (adj-*P*=1.4×10^-12^), the KstR1 and KstR2 regulons involved in cholesterol catabolism^50^ (adj-*P*=7.2×10^-5^ and 3.9×10^-4^, respectively), mycobactin genes that respond to iron scarcity^51^ (adj-*P*=6.4×10^-4^) and other processes (Online Analysis Tool). SEARCH-TB recapitulated specific previously described adaptations for persistence *in vivo,* including higher expression in untreated mice compared to the *in vitro* control of *icl1* (adj-*P*=3.8×10^-13^), the first gene of the glyoxylate bypass used during catabolism of fatty acids^52^, and increased expression of *tgs1* (adj-*P*=8.1×10^-35^) involved in triacylglycerol synthesis under environmental stress^53^.

**Fig. 3.**
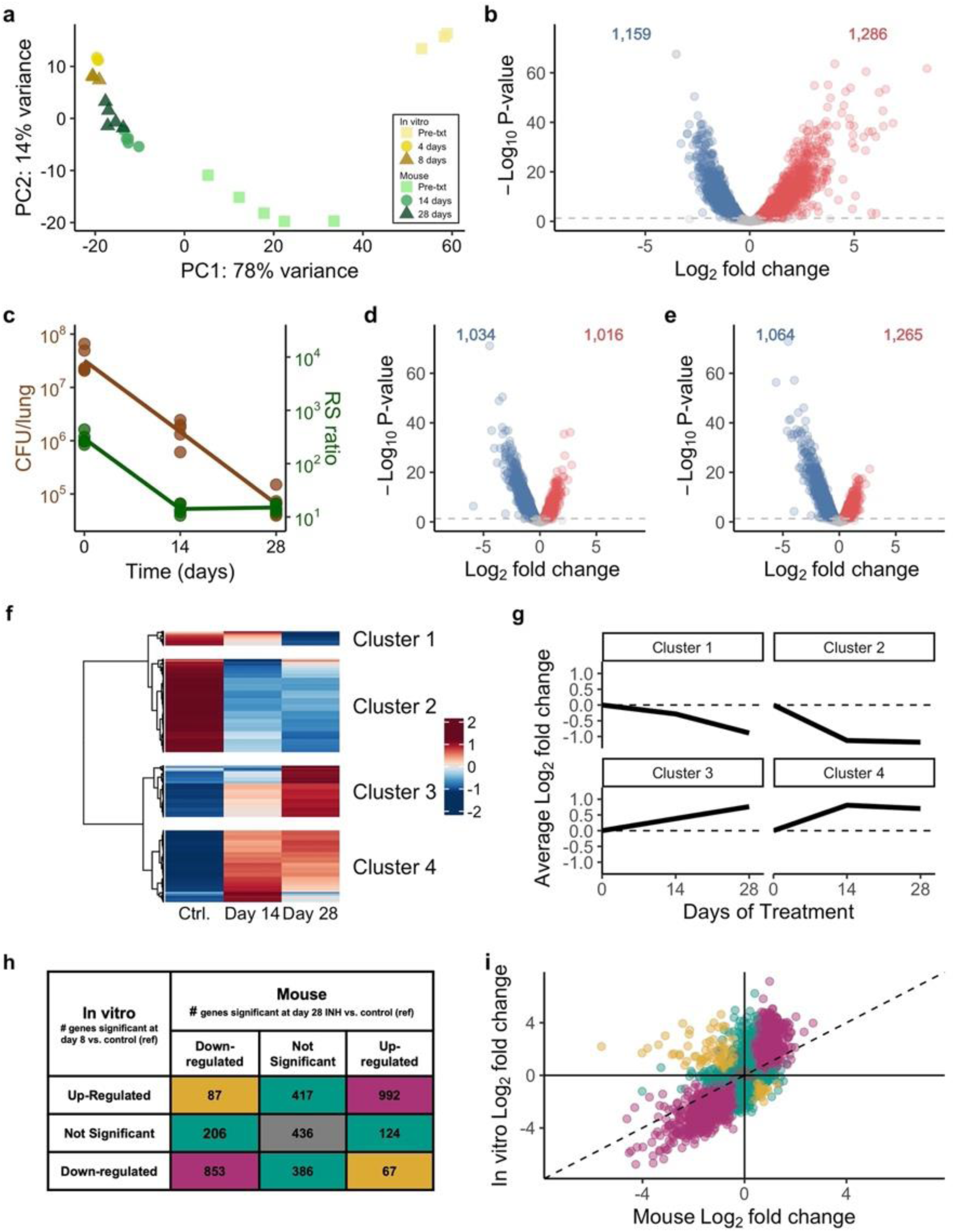
Overview of transcriptional response to HRZE in mice and *in vitro* experiments. **a.** Principal components plot including all mouse (triangles) and *in vitro* (squares) samples. Time points are shown by color. Volcano plot summarizing the differential expression between *Mtb* in mice and *in vitro* prior to treatment. Genes significantly down-(blue) or upregulated (red) in mice relative to *in vitro* (adj-*P*-value< 0.05) are shown. *Mtb* CFU and RS ratio values for control mouse samples and 14- or 28-days after HRZE treatment initiation. **d-e.** Volcano plots summarizing the differential expression between *Mtb* in 14-day HRZE treated and control mouse samples (**d**) and 28-day HRZE treated and control mouse samples (**e**). **f.** Estimated gene expression over time in mice. Genes that were significantly differentially expressed between at least two treatment time points are shown (N=2,429). Values are row-scaled, with red and blue indicating higher and lower expression, respectively. Hierarchical clustering of genes identified four broad patterns. **g.** Average log2 fold change for each of the four clusters relative to control. Values above and below zero represent up- and downregulation relative to control, respectively. **h.** Comparison of differential expression between mouse (day 28) or *in vitro* (day 8) relative to respective controls. Purple shading indicates genes with concordant fold-change direction and significance between mouse and *in vitro* experiments. Green shading indicates genes that were significant for either mouse or *in vitro* experiments but not both. Gold shading indicates genes that were significant for both mouse and *in vitro* experiments but in opposite directions. Gray shading indicates the genes that were not differentially expressed with HRZE treatment either in mouse or *in vitro* experiments. **i.** Comparison of fold-changes between mouse (day 28) or *in vitro* (day 8) relative to respective controls. Purple, green, gold, and gray colors have the same meaning as in **h**.

### Efficacy of drug treatment

In mice, treatment with HRZE reduced the CFU burden by 99.8% by day 28 (7.63 to 4.82 log_10_ CFU). A newer marker, the RS ratio,^10^ declined more rapidly than CFU, indicating that interruption of rRNA synthesis occurs more rapidly than killing (**Fig. 3c**).

### Global effect of HRZE on the *Mtb* transcriptome

In mice, treatment with HRZE transformed the *Mtb* transcriptome, significantly altering expression of 2,049 (57%) and 2,329 (65%) genes at day 14 and day 28 relative to pre-treatment control on day 0, respectively (**Fig. 3d-e**). A smaller set of genes (159 (4%)) was differentially expressed at day 28 relative to day 14. Hierarchical clustering identified clusters of genes with similar expression changes over time (**Fig. 3f**). The greatest transcriptional change occurred by day 14 (**Fig. 3g**, Clusters 2 and 4). A smaller subset of genes changed more gradually (**Fig. 3g**, Clusters 1 and 3). *In vitro* HRZE exposure also had a large impact on the *Mtb* transcriptome, altering expression of 2,748 (77%) and 2,800 (78%) of genes at day 4 and day 8 relative to control, respectively.

There were broad similarities as well as important differences between the effects of HRZE in mice and *in vitro*. Considering the effect of HRZE on gene expression at the latest treatment time points (8 days *in vitro* and 28 days in mice), differentially expressed genes were largely concordant between the mouse and *in vitro* samples (**Fig 3h**). However, eight days of HRZE *in vitro* induced overall larger fold-changes than 28 days of HRZE in mice (**Fig. 3i**). A notable subset of genes changed in discordant directions (colored gold in **Fig. 3i**), indicating differences between the effect of HRZE in mice and *in vitro* that are further described below.

### Effect of HRZE on bacterial cellular processes

Since our primary focus is *Mtb* phenotypes that survive prolonged drug exposure, the remainder of this manuscript describes only expression changes at the latest time point (28 days in mice or 8 days *in vitro*) relative to control, unless otherwise noted. Differences between other time points can be evaluated via the Online Analysis Tool.

Genes significantly impacted by 28-day HRZE treatment in mice were enriched for 36 of the 124 functional gene sets (**Table S3**) evaluated in enrichment analysis. Figure 4a provides a global portrait of change in cellular processes.

**Fig. 4.**
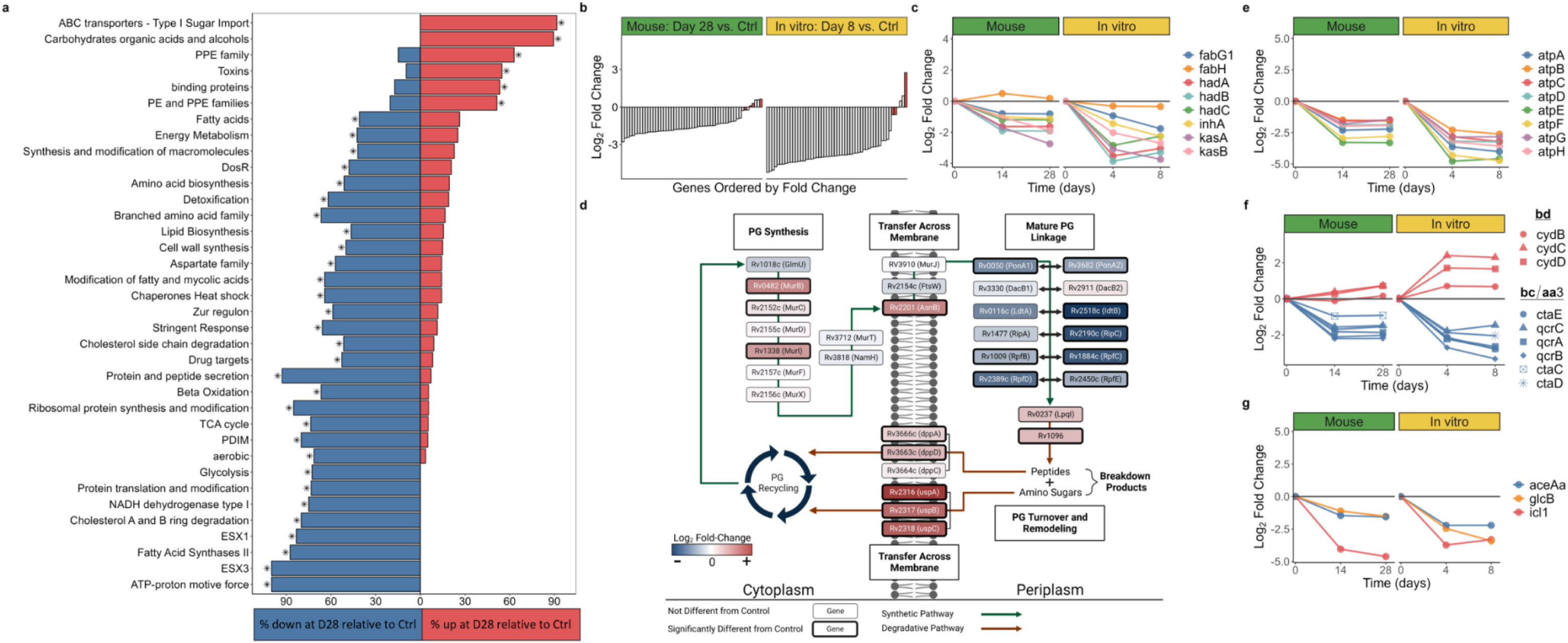
Summary of gene set enrichment and transcriptional changes in biological processes. **a.** Gene categories significantly enriched for genes differentially expressed between day 28 and control murine samples. The percentage of genes in each category significantly up- (red) or down- (blue) regulated for each comparison is illustrated. Asterisks indicate statistical significance (adj-*P*<0.05). **b.** Fold-change of ribosomal protein genes in mice at day 28 (left) and *in vitro* (right) at day 8, relative to control. Red bars indicate the four alternative C-ribosomal protein paralogs. **c, e-g** Fold-change values in mice at days 14 and 28 (left) and *in vitro* (right) at days 4 and 8, relative to control for FAS-II (**c**), ATP synthetase (**e**), cytochrome bcc/aa3 supercomplex and the bd oxidase (**f**), and glyoxylate bypass (**g**) gene sets. **d.** Graphical representation of changes in the peptidoglycan synthesis, modification, and recycling pathways in mice at day 28 relative to control. Log2 fold-change values for genes in the process are indicated by the color of each box and the bolded outlines of boxes represent genes that are significantly differentially expressed. Figure adapted from Maitra et al., 2019.^76^ Image created with Biorender.com.

#### Growth and synthesis of macromolecules

A dominant effect of HRZE was decreased expression of categories associated with growth and synthesis of macromolecules. Genes coding for ribosomal proteins were profoundly suppressed after treatment in mice (adj-*P*=2.9×10^-15^) and *in vitro* (adj-*P*=1.0×10^-14^) relative to their respective controls (**Fig. 4b**), consistent with decreased ribosome synthesis, a process fundamentally coupled with bacterial replication.^54, 55^ An exception to this pattern were the four alternative C-ribosomal protein paralogs lacking the zinc-binding CXXC motif that are a component of ribosomal remodeling. As shown in red in Figure 4b, genes for the alternative ribosomal proteins (*rpmB1, rpsR2, rpsN2, rpmG1*) had either sustained or significantly increased expression relative to control, consistent with ribosomal remodeling by a slowly replicating *Mtb* population.

Reduced protein synthesis was additionally suggested by decreased expression of the protein translation and modification category that includes genes responsible for translational initiation, promotion of tRNA binding, elongation, termination, and protein folding (adj-*P*=0.006 in mice, adj-*P=*0.007 *in vitro*). Concordant with downregulation of translation, transcription appeared suppressed, as evidenced by significant decrease in genes coding for gyrase A and B subunits (*gyrA, gyrB*) in mice and *in vitro* (least significant adj-*P=*1.8×10^-5^).

#### Cell-wall synthesis, remodeling, and recycling

##### Mycolic acids

Slowing of the initial step of mycolic acid synthesis was suggested by decreased expression of Rv2524c (*fas),* the gene coding for fatty acid synthetase I (FAS-I), in mice (adj-*P*=1.7×10^-9^) and *in vitro* (adj-*P*=3.2×10^-9^) and by decreased expression of the *fas* transcriptional regulator Rv3208^56^ in mice (adj-*P*=1.9×10^-4^) and *in vitro* (adj-*P=*1.1×10^-11^). Genes coding for the second step of elongation of acyl-coenzyme A to long chain fatty acids by fatty acid synthetase II (FAS-II) were also significantly decreased in both mice (adj-*P*=0.011) and *in vitro* (adj-*P=*0.027) (**Fig. 4c**). Finally, in *Mtb* treated with HRZE in mice and *in vitro,* there was decreased expression of the set of genes associated with elongation, desaturation, modification, and transport of the mature mycolic acids to the cell wall.

##### Phthiocerol dimycocerosates (PDIM)

Genes coding for PDIM biosynthesis, the outer surface glycolipids that are important for intracellular survival and virulence, were significantly downregulated in mice (adj-*P=*9.9×10^-5^) and *in vitro* (adj-*P=*0.017).

##### Peptidoglycan

In contrast to the downregulation observed in mycolic acid and PDIM genes, key genes involved in peptidoglycan synthesis, modification, and recycling had increased expression. Specifically, all five peptidoglycan synthesis genes assayed in the division cell wall operon (*ftsQ, murC, ftsW, murD, murR, murE*) had significantly increased expression at 14 days of HRZE and three remained significantly upregulated at day 28 after HRZE treatment in mice (Online Analysis Tool). Active recycling was also suggested by significantly increased expression with HRZE treatment in mice and *in vitro* of genes coding for the UspABC amino-sugar importer and the DppABCD dipeptide importer that transport peptidoglycan breakdown products (**Fig. 4d**).^57^

##### Trehalose

The TreY/Z genes that synthesize free trehalose from glycogen were significantly upregulated at all post-treatment time points in mice and *in vitro.* The other two trehalose synthesis pathways (OtsA/B and GlgE/TreS) were not differentially expressed with HRZE treatment in mice or *in vitro.* Remodeling of the trehalose component of the cell envelope was suggested by significantly increased expression of Rv3451 (*cut3*) under the test conditions in mice (adj-*P=*0.002) and *in vitro* (adj-*P=*4.2×10^-4^), which codes for a stress-responsive trehalose dimycolate hydrolase.^58^ Additionally, both mice and *in vitro* samples had significantly increased expression of four of the five genes for the LpqY/SugABC importer that is specific for the transport of trehalose.^59^

#### Metabolic readjustment

##### Electron transport and aerobic respiration

The eight genes that collectively code for ATP synthetase were strongly suppressed in mice (adj-*P*=7.6×10^-4^) and *in vitro* (adj-*P*=.003) (**Fig. 4e**). Oxidative phosphorylation appeared to transition from the primary cytochrome *bcc/aa3* supercomplex (downregulated) to the alternative less efficient cytochrome *bd* oxidase (upregulated) that has been implicated in persistence under environmental and drug stress^60^ (**Fig 4f**). Genes of the TCA cycle were downregulated with HRZE treatment in mice (adj-*P*=0.001) and *in vitro* (adj-*P=*0.010). Genes coding for NADH dehydrogenase types I and II and succinate dehydrogenase types I and II were also downregulated (Online Analysis Tool). By contrast, three of the four fumarate reductase genes were significantly induced by HRZE in mice, and all were induced *in vitro*.

##### Central carbon metabolism

Slowing of the TCA cycle was not accompanied with increased expression of glyoxylate bypass genes, an alternative pathway previously implicated in drug tolerance.^52^ Instead, the gene for isocitrate lyase (*icl1*), the first step of the glyoxylate bypass, was among the most profoundly suppressed by HRZE both in mice (adj-*P*=1.9×10^-37^) and *in vitro* (adj-*P*=2.0×10^-14^) (**Fig. 4g**). The gene coding for the alternative isocitrate lyase (*aceAa*) was also significantly downregulated during HRZE treatment in mice (adj-*P*=7.3.×10^-13^) and *in vitro* (adj-*P*=3.×10^-15^). Genes associated with carbon storage as triacylglycerol had discordant regulation in mice and *in vitro.* Specifically, *tgs1,* a gene in the DosR regulon which codes for triacylglycerol synthase, decreased 4.4-fold (adj-P=7.0×10^-13^) in mice but increased 10.9-fold (adj-P=3.2×10^-16^) *in vitro*.

#### Cholesterol degradation

Although more highly expressed in mice than *in vitro* prior to drug exposure, genes related to cholesterol breakdown were suppressed by HRZE in mice and increased by HRZE *in vitro* (**Fig 5a**, significance of individual genes in Online Analysis Tool).

**Fig. 5.**
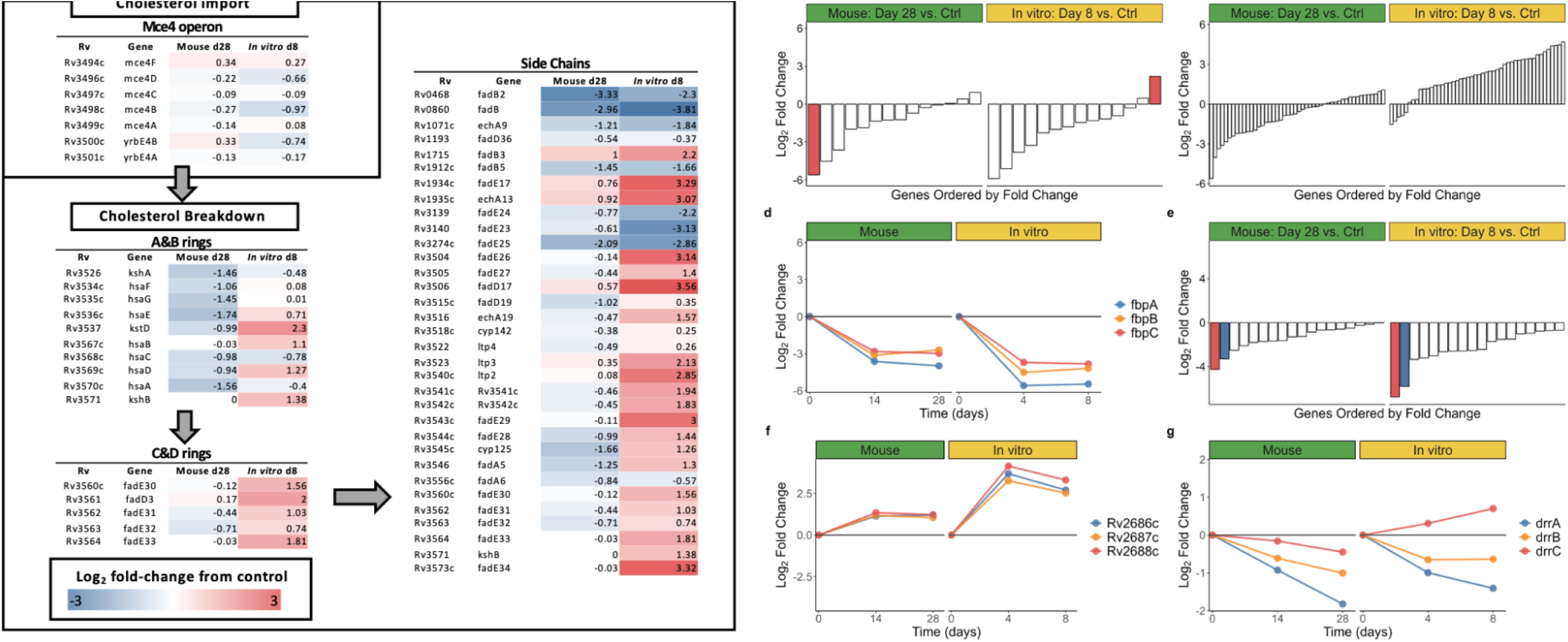
Summary of transcriptional changes in biological processes. **a.** Log2 fold-change values for genes in the cholesterol degradation pathway for mice at day 28 and *in vitro* at day 8, relative to control. Deeper red values represent higher upregulation of genes after drug treatment relative to control, while deeper blue values represent higher downregulation. Figure adapted from Pawełczyk et al., 2021^77^. **b-g.** Fold-change in mice (left) and *in vitro* (right), relative to control for heat shock proteins (red bar represents *hspX*) (**b**), dosR (**c**), antigen 85 (**d**), ESX-1 (red bar represents *esxA*, blue bar represents *esxB*) (**e**), the Rv2686c-2688c efflux pump (**f**), and the DrrABC efflux pump (**g**) gene sets.

##### Glycerophosphodiesters

The four genes coding for the UgpABCE transporter responsible for import of glycerophosphodiesters that are a major source of carbon and phosphate^61^ were strongly upregulated at both treatment time points in mice and *in vitro* (Online Analysis Tool).

#### Lipolytic enzymes

The complex repertoire of lipolytic enzymes that are implicated in *Mtb* dormancy and virulence includes Lipase family genes predicted based on sequences to have lipase/esterase activity. In mice, 11 of the lipase genes included in the SEARCH-TB assay had significantly increased expression, including *lipY* that codes for triacylglycerol lipase LipY. There was particularly strong induction of *lipY in vitro* (15.5-fold increase, *P=*1.2×10^-35^). Additionally, three of the four phospholipase C genes (*plcA, plcB, plcC*) were significantly upregulated in mice.

#### Canonical stress responses

Although the 14 heat shock proteins (HSPs) that act as chaperones in protein folding are often described as a stress response, expression of HSP genes largely decreased following HRZE exposure. A notable discordance between mice and *in vitro* results is the hypoxia-responsive *hspX* (*arc*) (noted in red in **Fig. 5b**) that had the greatest negative fold-change of all genes evaluated in mice (48.8-fold decrease, adj-*P=*4.8×10^-57^) yet was significantly upregulated *in vitro* (4.6-fold increase, adj-*P*=2.0×10^-8^). Concordant with the *hspX* result, we found that HRZE significantly decreased expression of the DosR regulon in mice (adj-*P=*0.032) but induced the DosR regulon *in vitro* (adj-*P*=2.2×10^-7^) (**Fig. 5c**). Genes of the stringent response, which is typically induced by nutrient deprivation and other environmental stresses, were significantly downregulated with HRZE treatment in mice (adj-*P=*2.9×10^-15^) and *in vitro* (adj-*P=*1.6×10^-14^) (Online Analysis Tool). Similarly, *relA,* the gene coding for the stringent response regulator was significantly decreased in mice (adj-*P*=2.0×10^-6^) and *in vitro* (adj-*P=*7.0×10^-6^).

#### Secretion of peptides and immunogenic proteins

Genes coding for the Antigen 85 complex, the major secreted protein that is essential for intracellular survival within macrophages^62^, were profoundly downregulated under the test conditions in mice (smallest fold-decrease=6.5, least-significant adj-*P=*3.1×10^-24^) and *in vitro* (smallest fold-decrease=14.2, least-significant adj-*P=*2.6×10^-31^) (**Fig. 5d**). Expression of the ESX-1 locus was downregulated (adj-*P=*9.4×10^-5^ in mice, adj-*P=*1.3×10^-5^ *in vitro*) with particularly strong suppression of *esxA* and *esxB,* genes coding for highly-immunogenic early secretory antigenic 6 kDa (ESAT-6) and culture filtrate protein 10 (CFP-10), in mice (smallest fold-decrease=10.0, least-significant adj-*P=*4.7×10^-24^) and *in vitro* (smallest fold-decrease=55.7, least-significant adj-*P=*2.4×10^-35^) (**Fig. 5e**). Expression of all 9 genes assayed in the ESX-3 locus was significantly downregulated with particularly strong suppression of immunogenic *esxG* and *esxH* in mice (smallest fold-decrease=18.5, least-significant adj-*P=*1.1×10^-34^) and *in vitro* (smallest fold-decrease=44.3, least-significant adj-*P=*4.0×10^-32^). In contrast, six of the seven genes in the ESX-4 locus that a recent review^63^ described as having “wholly unknown” function had significantly increased expression. Interestingly, the Sec-independent Tat system that exports pre-folded proteins had significantly increased expression of genes coding for the TatBC complex (*tatB, tatC*) but significantly decreased expression of the gene for the PMF-dependent TatA pore protein (*tatA*).

#### Efflux pumps

Most of the diverse set of transporters that includes putative drug efflux pumps had altered expression after HRZE exposure, but the direction of their regulation varied (**Table S4**). As examples, Rv2686c-2688c that is associated with fluoroquinolone tolerance^64^ was upregulated in mice and *in vitro* (**Fig. 5f**) but DrrABC that is associated with daunorubicin tolerance and appears in clinical drug-resistant strains^64^ was downregulated (**Fig. 5g**).

#### Transcriptional and post-transcriptional regulation

SEARCH-TB indicated large-scale regulatory reprogramming. For example, most of the 188 transcription factors assayed in mice changed after 28 days of HRZE treatment, with 66 significantly increased and 47 significantly decreased. The regulatory perturbation was even more pronounced *in vitro* with 87 significantly increased and 55 significantly decreased transcription factors after 8 days of HRZE treatment. Of the 12 sigma factor genes included in SEARCH-TB, *sigF, sigI* and *sigM* were significantly upregulated following HRZE treatment in both mice and *in vitro. sigA, sigB, sigD* and *sigK* were significantly downregulated in both mice and *in vitro.* Expression of *sigC* and *sigL* was significant in both mice and *in vitro* but in discordant directions, suggesting differing regulatory responses *in vivo* and *in vitro*.

HRZE appeared to activate the post-transcriptional toxin-antitoxin system that modulates the concentration of existing transcripts. Toxin genes had significantly increased expression in mice (adj-*P=*0.011) and *in vitro* (adj-*P=*0.027). The counter-regulatory antitoxins that restrict toxin activity were not categorically altered in mice but were significantly suppressed *in vitro* (adj-*P*=0.037).

#### Drug targets

Because processes that are upregulated following prolonged drug exposure may represent survival mechanisms that could be targeted to eradicate persisting *Mtb,* we evaluated genes that code for drug targets. Evaluation of targets of 31 existing drugs or investigational compounds (36 genes) (**Table S5**) were predominantly suppressed in mice (19 down-regulated and 3 upregulated) and *in vitro* (26 down-regulated and 2 upregulated). Notable exceptions to downregulation of drug targets were increased expression of the genes for Mur ligases B and C that initiate peptidoglycan synthesis in mice^65^ and increased expression of *rfe,* the gene for phosphoglycosyltransferases WecA that initiates arabinogalactan synthesis in mice and *in vitro.*^66^

### Transcriptional differences between *Mtb* in mice and *in vitro* at final time points

After 8 and 28 days of HRZE treatment, the *in vitro* and murine *Mtb* transcriptomes were more similar than they were prior to HRZE exposure (**Fig 3a**). Nonetheless, 1,014 genes (28%) remained differentially expressed between the final murine and *in vitro* time points.

## DISCUSSION

This work established the SEARCH-TB platform, a method for elucidating the *Mtb* transcriptome during drug exposure *in vivo*. After 28 days of treatment with the global standard 4-drug combination (HRZE) and 99.8% reduction in the burden of culturable *Mtb* in mouse lungs, SEARCH-TB indicated broad suppression of cellular activity including slowing of metabolism, synthesis of macromolecules and secretion of immune-modulating peptides. SEARCH-TB also suggested bacterial adaptation to drug stress, including a shift in electron transport to the alternative less efficient cytochrome *bd* oxidase, ribosomal remodeling, cell wall remodeling and recycling, and reprogramming of regulatory and efflux pump activity. The effects of HRZE in mice and *in vitro* had both broad similarities and notable differences that likely reflect effects of immunity and environment. As the first platform capable of nearly genome-wide quantification of extremely low abundance *Mtb* transcripts in mice, SEARCH-TB should enable highly granular evaluation of the effect of drugs and regimens *in vivo*.

Unsurprisingly, at baseline, prior to the start of HRZE treatment, SEARCH-TB indicated that *Mtb* cellular processes differed substantially between mice and early log phase growth, reflecting bacterial adaptation to immunity and the lung environment. These differences between untreated mouse and *in vitro* control are consistent with a recent review^47^ of the treatment-naïve *in vivo Mtb* transcriptome, both in terms of broad cellular processes (downregulation of genes associated with transcription, translation and metabolism *in vivo*) and specific adaptations (*e.g.,* increased expression of genes of the DosR regulon, glyoxylate bypass genes). Importantly, the transcriptome of the untreated mouse was the starting point for our current analysis of drug effect. Combination antimicrobial treatment is a categorically different type of stress than environmental conditions such as pH, hypoxia, and nutrient starvation. Correspondingly, as highlighted in the discussion below, SEARCH-TB showed that drug exposure elicited transcriptional adaptations that often diverge from well-established transcriptional responses to environmental stress.

SEARCH-TB results indicated that a major effect of HRZE treatment in mice is suppression of *Mtb* growth and metabolism. Consistent with previous *in vitro* analyses of drug effects, transcription results suggested decreased synthesis of all major macromolecules, including rRNA, protein, lipids, and cell wall constituents. An additional manifestation of decreased bacterial activity was decreased protein and peptide secretion, including the ESX1 Type VII secretion system that exports the highly immunogenic ESAT-6 and CFP-10 proteins. This suggests that drug stress might alter the pathogen’s capacity to modulate host immunity.

Despite the broad downregulation of activity, SEARCH-TB provided evidence that drug-stressed *Mtb* is not inert or incapacitated. Increased expression of genes associated with peptidoglycan and trehalose synthesis and recycling suggested active cell wall modification. SEARCH-TB indicated broad transcriptional and post-transcriptional regulatory reprogramming and reconfiguration of efflux pump expression. HRZE appears to have induced reconfiguration of metabolism and energy generation away from high respiratory activity that maximizes ATP generation, and towards reduced respiratory efficiency and oxidative phosphorylation activity.

A striking finding was that HRZE led to significantly decreased expression of the DosR regulon in mice. DosR is the *Mtb* response to impaired aerobic respiration induced by hypoxia, nitric oxide, or carbon monoxide exposure.^67^ DosR was termed the “dormancy survival regulon” because impaired respiration leads to growth arrest. In mice, HRZE appeared to arrest growth yet expression of the DosR regulon was suppressed. We hypothesize that once 99.8% of culturable *Mtb* have been eliminated and secretion of immunogenic peptides, including Ag85, ESAT-6 and CFP-10, is diminished in the residual population, host immunity may be modulated. A less-intense inflammatory response would be predicted to decrease increase localized oxygen availability and decrease nitric oxide production, thereby diminishing induction of DosR-inducing conditions that restrict aerobic respiration. The hypothesis that DosR may be an indirect readout of immune response is consistent with our previous observation that the DosR regulation has significantly lower expression in TB patients with AIDS than in immunocompetent patients.^33^ Also consistent with this hypothesis was our finding that *in vitro*, in the absence of immunity, HRZE significantly increased rather than decreased expression of DosR genes.

Also notable was expression of the gene for isocitrate lyase, the first step of the glyoxylate bypass that is essential to establishing infection in animals.^68–70^ Consistent with these prior results, we found that *icl1* was highly expressed in untreated mice relative to log phase growth. *icl1* was also shown to have increased expression following sublethal exposure to rifampin, INH or streptomycin *in vitro* and was identified as a mediator of drug tolerance.^52^ By contrast, we found that treatment with lethal doses of HRZE strongly suppressed, rather than induced, expression of both *icl1* and *aceAa* which codes for an alternative isocitrate lyase. This highlights, first, that adaptations associated with prolonged tolerance to HRZE differ from adaptations to host environments that enable persistent infection in the absence of drug therapy and, second, that the response to exposure to a combination regimen *in vivo* may differ from single drug exposure *in vivo*.

Drugs have historically targeted transcription, translation, and other growth-associated cellular processes^71^ that SEARCH-TB shows are downregulated after treatment with HRZE. While transcriptional downregulation of drug targets does not necessarily mean a drug will be ineffective, genes such as the Mur ligases that have increased expression after 1-month of HRZE treatment *in vitro* are noteworthy as they might indicate adaptations that enable *Mtb* to withstand drug exposure. Similarly, genes coding for the alternative bd oxidase that is a proposed drug target^72^ were upregulated at a time when nearly all other metabolism-associated genes had suppressed expression.

A novel comparison in this report is the effect of HRZE in the mouse versus *in vitro.* Although the physiologic state of *Mtb* appeared quite different in mice and *in vitro* at pre-treatment baseline, HRZE induced broadly similar changes *in vivo* and *in vitro*. The magnitude of fold-change was greater *in vitro,* potentially indicating a lower effective drug exposure in the mouse. Nonetheless, we also observed discordant transcriptional changes such as in expression of the DosR regulon, indicating that drug effect *in vitro* is not a simple surrogate for effect in the mouse. Furthermore, at the latest treatment endpoints, a large number of genes remained differentially expressed between *in vitro* and the mouse, likely indicating the importance of immunity and pharmacokinetics in the mouse.

As a highly granular readout of the effect of TB treatment on bacterial cellular processes, SEARCH-TB may improve precision of pharmacodynamic evaluation, providing greater information than the existing standard method of assessing drug effectiveness *in vivo* (*i.e.,* culture-based enumeration of bacterial burden^73^). We have previously shown that CFU burden does not capture the entirety of complex drug effects *in vivo*. For example, regimens that have identical effects on CFU in mice can have different long-term relapse outcomes.^10, 74^ In previous studies, we demonstrated that quantification of rRNA synthesis via the RS ratio showed proof of concept that molecular measures of bacterial cellular processes *in vivo* can distinguish regimens that are indistinguishable based on CFU.^10, 74^ SEARCH-TB advances molecular characterization of drug effects *in vivo* to a higher level of granularity. Others have demonstrated the power of *Mtb* transcriptional readouts of drug effect to predict drug interactions *in vitro.*^75^ We believe that SEARCH-TB has potential to enable a new era of pharmacodynamic evaluation in which drug interactions and regimens are assessed based on molecular effects on cellular process *in vivo*.

This report has several limitations. First, between-target variation in the amplification efficiency of SEARCH-TB primers likely affects the rank-order of gene counts, meaning that, for individual genes, a higher absolute count does not necessarily indicate greater expression. However, because amplification is highly repeatable, the modest amplification bias identified does not affect estimation of differential expression between groups which is the primary purpose of SEARCH-TB. Indeed, we showed that SEARCH-TB provides the same biological interpretation as conventional RNA-seq. Second, specific and efficient primers could not be designed for ∼12% of *Mtb* transcripts. Third, transcriptional profiling is inherently unable to resolve the enduring question of whether the drug-tolerant subpopulation that persists late into treatment results from selection (*i.e.,* elimination of easily-killed *Mtb*) or physiological transformation of *Mtb* that were present at baseline. Finally, this initial demonstration of SEARCH-TB evaluated a single regimen (HRZE) in a single murine model. Important next steps include evaluation of individual drugs and diverse regimens in additional animal models.

SEARCH-TB enabled what to our knowledge is the first evaluation of the effect of prolonged drug treatment on *Mtb* transcription in animal models, revealing adaptations distinct from those observed under environmental stress. The *Mtb* subpopulation that survived one month of HRZE treatment appeared substantially less active than prior to treatment but was not inert with transcriptional changes suggesting adaptation for survival. SEARCH-TB should enable a new era of *in vivo* molecular pharmacodynamics with potential to accelerate identification of new highly potent regimens.

**Data availability**. All raw sequencing data have been deposited in the Sequence Read Archive (SRA) under BioProject accession PRJNA939248. Individual samples have BioSample accession numbers SAMN33461189 through SAMN33461251.

**Funding**. GR acknowledges funding from the Bill and Melinda Gates Foundation (INV-009105). NDW and RD acknowledge funding from the Bill and Melinda Gates Foundation (OPP1170003). NDW and PN acknowledge funding from the US National Institutes of Health (1R01AI127300-01A1). NDW acknowledges funding from Veterans Affairs (1I01BX004527-01A1).

## Acknowledgements

We are grateful to the Illumina Design Team that developed the custom primer pool used in SEARCH-TB.

