## Supplemental Information for "Transcriptional adaptation of drug-tolerant *Mycobacterium tuberculosis* in mice"

### 1. **Supplemental Methods**

#### 1.1. ***Table S1****. Inclusion reference genomes.*

Genomes from eight *M tuberculosis* Complex strains used in the design process to assure that SEARCH-TB amplifies sequence that is common across lineages.

| Species | Strain Name | Lineage | Genbank accession number |
| --- | --- | --- | --- |
| *Mycobacterium tuberculosis* | EAI5 | L1 | NC_021740 |
| *Mycobacterium tuberculosis* | CCDC5180 | L2 | NC_017522 |
| *Mycobacterium tuberculosis* | CAS/NITR204 | L3 | NC_021193 |
| *Mycobacterium tuberculosis* | CDC1551 | L4 | NC_002755 |
| *Mycobacterium tuberculosis* | H37Rv | L4 | NC_000962 |
| *Mycobacterium tuberculosis* | Erdman | L4 | AP012340.1 |
| *Mycobacterium africanum* | GM041182 | L6 | NC_015758 |
| *Mycobacterium bovis* BCG | Pasteur1173P2 | L8 | NC_008769 |

#### 1.2. ***Table S2****. Exclusion reference genomes*

Genomes from 12 eukaryotes and bacteria used in the design process to assure that SEARCH-TB does not amplify sequence from non-*M tuberculosis* complex species.

| Species | Strain Name |
| --- | --- |
| *Homo sapiens* |  |
| *Mus musculus* | C57BL/6J |
| *Mycobacterium avium* | H87 |
| *Mycobacterium abscessus* | ATCC19977 |
| *Mycobacterium kansasii* | ATCC12478 |
| *Escherichia coli* | K-12 |
| *Streptococcus mutans* | UA159 |
| *Rhodococcus equi* | 103S |
| *Pseudomonas aeruginosa* | PAO1 |
| *Prevotella intermedia* | 17 |
| *Neisseria bacilliformis* | AFAY01 |
| *Nocardia asteroids* | FOTX01 |

#### 1.3. ***Conversion to Mtb Erdman***

*Mtb* H37Rv genes targeted by the SEARCH-TB panel were mapped to the Erdman strain and genes without an adequate match were removed from further analysis. The Basic Local Alignment Search Tool (BLAST)^1^ was used to find matches between the panel primer sequences and the Erdman genome (annotated genome obtained from RefSeq; annotation date: 06/07/2020). Genes were excluded from the panel if their corresponding primers did not map to a gene in the Erdman genome, there were more than 2 mismatches between either primer and the best match to the Erdman genome, or the location of the matches in the Erdman genome for a set of primers were more than 160 base pairs apart.

#### 1.4. ***Evaluation of amplification bias***

We tested for amplification bias (*i.e.,* differences in amplification efficiency between primer pairs targeting different *Mtb* transcripts) by sequencing *Mtb* genomic DNA with SEARCH-TB. The value of DNA for this purpose is that each gene is present as a single copy in the genome. To avoid overloading the amplification steps with template material, 0.001 ng *Mtb* H37Rv genomic DNA was diluted in water and sequenced in duplicate using SEARCH-TB. The raw counts for each sample were transformed to counts per million (CPM). In the absence of amplification bias (*i.e.,* if all primer amplified with identical efficiency), the expected CPM would be one million divided by the number of targets. We compared observed CPM for each target with the ideal CPM expected in the absence of bias.

#### 1.5. ***Evaluation of repeatability of amplification***

The repeatability of amplification was evaluated by evaluating samples with a known amount of *Mtb* RNA. To prepare these samples, we spiked 0.001 ng *Mtb* RNA, collected at log-phase growth into 1 ng human lung RNA purchased from Invitrogen. Aliquots of this spiked-in mixture were sequenced with SEARCH-TB.

#### 1.6. ***Concordance of SEARCH-TB with conventional RNA-seq***

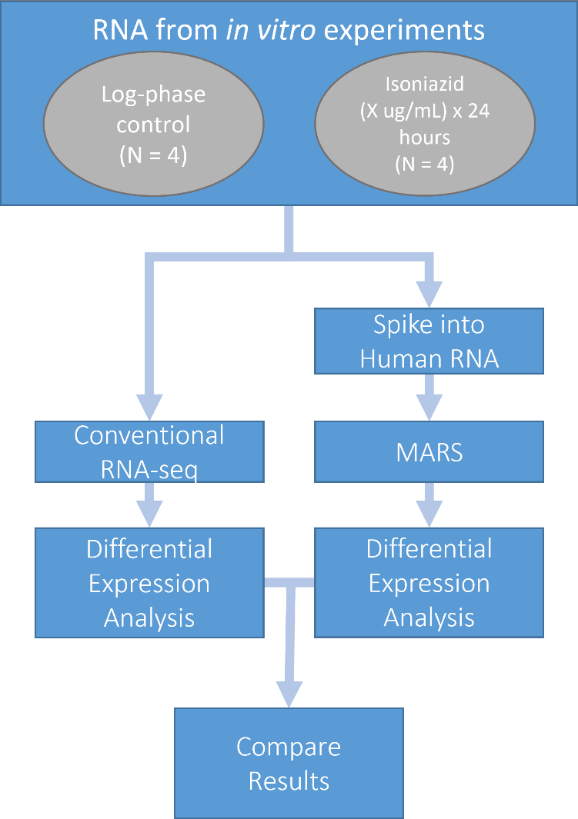
Our purpose was to determine if SEARCH-TB provided the same biological information that would be obtained by conventional RNA-seq (**Fig S1**). *Mtb* was grown to early log phase and exposed to isoniazid (0.1 ug/mL) for 24 hours. RNA was extracted from control and isoniazid-treated samples. Control and isoniazid-treated RNA aliquots (N=4 each) were prepped using the Illumina TrueSeq protocol and sequenced on the NovaSeq as described in Sections 1.10-1.11 below. We described this sequencing strategy which did not include *Mtb-*targeted amplification as our conventional RNA-seq reference. Additional control and isoniazid-treated RNA aliquots (N=4 each) were also spiked into human RNA at a ratio of 1:1,000. Libraries were prepped via SEARCH-TB and sequenced on the NovaSeq as described in Sections 1.10-1.11. For both the conventional RNA-seq and SEARCH-TB results, we calculated differential expression between the control and isoniazid-treated samples using edgeR. We evaluated the concordance of fold-changes and differentially expressed gene lists between conventional RNA-seq and SEARCH-TB.

**Fig. S1**. Conceptual diagram for comparison of SEARCH-TB with conventional RNA-seq.

#### 1.7. ***In vitro drug experiments***

For *in vitro* drug exposure, *Mtb* strains H37Rv and/or Erdman were exposed to 0.1 μg/ml isoniazid or HRZE with individual component concentrations of 0.1, 0.1, 3.2, and 4.0 μg/ml for isoniazid, rifampin, pyrazinamide, and ethambutol respectively. Drug exposure experiments were carried out as described in Walter et al., 2021.^2^ Cultures were collected for RNA after 24 hour isoniazid exposure and 4 days and 8 days of HRZE exposure. Drugs were obtained from the following suppliers: isoniazid (Sigma, cat# I1377), rifampicin (Chem-Impex Int’l, cat# 00260), pyrazinamide (Sigma, cat# PHR1576), ethambutol (Sigma, PHR1930). Drug stocks were prepared in water (isoniazid, ethambutol) or DMSO (rifampicin, pyrazinamide) at 10 mg/ml, filter sterilized, and stored at -80^o^C. Drugs were diluted to 1000X working concentration immediately prior to use.

#### 1.8. ***Murine drug experiments***

Female BALB/c, 6- to 8-week old mice, were infected with *Mtb* Erdman by high dose aerosol using a GlasCol chamber as described previously (PMID: 34006838). Treatment started 11 days later (D0). Mice were sacrificed for lung CFU counts on the day after infection and D0 to determine the number of CFU implanted and the number present at the start of treatment, respectively. Lungs were dissected aseptically, and flash frozen in liquid nitrogen before processing. Rifampin (R) was administered at 10 mg/kg. Isoniazid (H) was administered at 10 mg/kg. Pyrazinamide (Z) was administered at 150 mg/kg. Ethambutol (E) was administered at 100 mg/kg. All drugs were prepared in sterile water and given by oral gavage 5 of 7 days per week. HZE was administered in combination 1 hour after mice received R. Assessment of bactericidal activity were based on lung CFU counts after 2 or 4 weeks of treatment. At each time point, lungs were dissected aseptically, and the left lung, and inferior and post-caval lobes (2/3rds of the total lung by mass) were homogenized in 4.5 ml PBS + 10% [w:v] bovine serum albumin (BSA). Lung homogenates were plated in serial dilutions on 0.4% [w:v] charcoal-supplemented 7H11 agar supplemented with 10% oleic acid, BSA, sodium chloride, dextrose and catalase (OADC) and with selective antibiotics: cycloheximide (10 mg/L), carbenicillin (50 mg/L). The remaining superior and middle lung lobes (1/3rd of the total lung by mass), were recovered and flash frozen under liquid nitrogen for RNA preservation.

#### 1.9. ***RNA extraction***

*In vitro* cells were collected in a 5M GTC-TCEP solution^3^ at a ratio of one volume culture to two volumes GTC-TCEP. Cultures were mixed and incubated at room temperature for five minutes. Mycobacterial cells were pelleted by centrifugation at 4000 rpm for five minutes. Cell pellets were resuspended in 1 mL RLT+ BME (Qiagen, 1% BME) with 0.5 mL 0.1 mm glass beads. Cells were lysed by beadbeating using a FastPrep-24 (MP Biomedical) at 4.0 m/s for 30 seconds four times, resting on ice for two minutes each time.

Murine tissue samples were snap frozen in 7 mL Precellys tubes (Bertin) under liquid nitrogen. To each tissue sample, 1.5 mL GTC-TCEP buffer^3^ was added and samples were homogenized at 7,200 rpm for 16s using a Precellys Evolution (Bertin). Homogenates were incubated at room temperature for 5 minutes while protected from light. Unlysed mycobacterial cells were pelleted at 10,000g for 3 minutes, then resuspended in 0.8 mL RLT+ BME buffer (Qiagen, TCEP concentration 0.75 mM). Mycobacterial cells were lysed on a Precelly Evolution (Bertin) at 6,500 rpm for 30s three times, resting on ice for 5 minutes each time, and stored at -80⁰C.

After storage at -80⁰C, *in vitro* and murine lysates were thawed and RNA was purified using the Maxwell simplyRNA cells kit (*in vitro* samples) or simplyRNA tissue kit (murine samples) using the Maxwell RSC instrument following the manufacturer’s instructions with the following modifications. Additional kit DNase was added at twice the manufacturer’s recommended amount. Purified RNA was quantified using the Quantifluor RNA system (Promega).

#### 1.10. ***Library preparation***

This project prepared two library types (SEARCH-TB and Illumina RNA-seq). This section will describe these two library preparations sequentially.

Preparation of SEARCH-TB libraries

Samples were prepared for SEARCH-TB using Illumina’s AmpliSeq for Illumina Custom and Community RNA Panels kit, according to the Illumina’s recommendations except where noted. RNA was diluted to a concentration of 15-30 ng/uL, and reverse transcribed. cDNA targets were amplified, with the SEARCH-TB panel at 2x concentration instead of 5x, with the following thermocycling conditions: 99°C for 2 minutes, 18 cycles of 99°C for 15 seconds and 60°C for 8 minutes, hold at 10°C. Amplicons were partially digested, indexes were ligated, and the library was cleaned. The library was amplified, with the following thermocycling conditions: 98°C for 2 minutes, 9 cycles of 98°C for 15 seconds and 64°C for 1 minute, hold at 10°C. The library was cleaned a second time, quantified via Qubit using the dsDNA kit (Invitrogen), and evaluated for quality via TapeStation with the High Sensitivity D1000 ScreenTape (Agilent). Libraries were diluted to the same concentration and checked for intact sequencing adaptors via qPCR. Inserts were 124-139 bp, yielding amplicons of approximately 280 bp.

RNA-seq libraries were prepared with 100 ng of high-RIN RNA using the Truseq Stranded Total RNA-seq library prep kit with RiboZero Plus as per protocol (Illumina). Briefly, samples were depleted of rRNA with the RiboZero Plus depletion kit, denatured and fragmented, followed by cDNA generation and adaptor ligation. Libraries were targeted for sequencing of 10 million paired-end 2x150 bp reads on a NovaSeq 6000 instrument (Illumina).

#### 1.11. ***Sequencing and bioinformatic analysis***

Libraries were sequenced on an Illumina NovaSeq6000 at the University of Colorado Anschutz Medical Campus Genomics Shared Resource, 2x150. 8,886,900 - 92,226,565 pairs of raw sequences were obtained per library. Bioinformatics analyses were performed on the Health Data Compass Eureka v1.0 at the University of Colorado Anschutz Medical Campus. Adaptors and bar codes were removed, and sequences were quality-trimmed with Skewer, with a minimum Qscore of 20 and a length between 50-175 bp.^4^ FastQC was used to confirm the size distribution and high quality of the remaining sequences. High-quality sequences were mapped to *Mtb* Erdman using Bowtie2 v2.4.0 using the default parameter.^5^ Mapped sequences were counted using HtSeq v1.0 with the default parameters.^6^

#### 1.12. ***Table S3.*** *Curated gene categories used for enrichment analysis*

Gene categories curated from the literature which were used in enrichment analysis. Along with the reference, the total number of genes from each source along with the number of genes from each source which were in the SEARCH-TB assay is given.

| **Category** | **# of Genes in Assayed** | **# of Genes in Category** | **Source** |
| --- | --- | --- | --- |
| ABC transporters - Type I Sugar Import | 12 | 12 | (Soni, Dubey, & Bhatnagar, 2020)^7^ |
| ABC transporters Type I phosphate | 8 | 8 | (Soni, Dubey, & Bhatnagar, 2020)^7^ |
| Antitoxins | 72 | 76 | (Shao et al., 2011)^8^ |
| Arabinogalactan (AG) | 18 | 19 | (Abrahams & Besra, 2018)^9^ |
| Beta Oxidation | 18 | 18 | (Schnappinger et al., 2003)^10^ |
| Cell wall synthesis | 40 | 40 | (Kirksey et al., 2011)^11^ |
| Cholesterol A and B ring degradation | 10 | 10 | (Pawełczyk et al., 2021)^12^ |
| Cholesterol side chain degradation | 33 | 33 | (Pawełczyk et al., 2021)^12^ |
| DNA replication and repair | 25 | 27 | (Ditse, Lamers, & Warner, 2017)^13^ |
| DosR | 48 | 48 | (Voskuil et al., 2003)^14^ |
| Drug targets | 40 | 40 | (“Working Group for New TB Drugs.,” 2021)^15^  (Shetye, Franzblau, & Cho, 2020)^16^ |
| Efflux Pumps and Transports | 25 | 26 | (Remm, Earp, Dick, Dartois, & Seeger, 2022)^17^ |
| Enduring Hypoxic Response | 149 | 161 | (Rustad, Harrell, Liao, & Sherman, 2008)^18^ |
| Esterases (Lip family) | 20 | 22 | (Tallman, Levine, & Beatty, 2016)^19^ |
| Esterases (non-Lip family) | 13 | 13 | (Tallman, Levine, & Beatty, 2016)^19^ |
| ESX1 | 18 | 19 | (Gröschel, Sayes, Simeone, Majlessi, & Brosch, 2016)^20^ |
| ESX2 | 12 | 12 | (Gröschel, Sayes, Simeone, Majlessi, & Brosch, 2016)^20^ |
| ESX3 | 9 | 11 | (Gröschel, Sayes, Simeone, Majlessi, & Brosch, 2016)^20^ |
| ESX5 | 11 | 15 | (Gröschel, Sayes, Simeone, Majlessi, & Brosch, 2016)^20^ |
| Fatty Acid Synthases II | 8 | 9 | (Duan, Xiang, & Xie, 2014)^21^ |
| kstR1 regulon | 70 | 71 | (Wipperman, Sampson, & Thomas, 2014)^22^ |
| kstR2 regulon | 14 | 15 | (Wipperman, Sampson, & Thomas, 2014)^22^ |
| LAM | 14 | 15 | (Batt, Burke, Moorey, & Besra, 2020)^23^ |
| mmpL | 14 | 14 | (Domenech, Reed, & Barry, 2005)^24^ |
| Mycobactin Biogenesis | 10 | 10 | (Quadri, Sello, Keating, Weinreb, & Walsh, 1998)^25^ |
| Mycolic acid modification | 12 | 12 | (Marrakchi, Lanéelle, & Daffé, 2014)^26^ |
| NADH dehydrogenase type I | 12 | 14 | (Cook, Hards, Vilchèze, Hartman, & Berney, 2014)^27^ |
| Oxidative Stress | 48 | 49 | (Voskuil, Bartek, Visconti, & Schoolnik, 2011)^28^ |
| PDIM | 20 | 20 | (Rens, Chao, Sexton, Tocheva, & Av-Gay, 2021)^29^ |
| Peptidoglycan (PG) | 32 | 34 | (Maitra et al., 2019)^30^ |
| Sigma Factors | 12 | 13 | (Lew, Kapopoulou, Jones, & Cole, 2011)^31^ |
| Stringent Response | 123 | 147 | (Dahl et al., 2003)^32^ |
| Toxin-Antitoxin | 146 | 152 | (Shao et al., 2011)^8^ |
| Toxins | 74 | 76 | (Shao et al., 2011)^8^ |
| Transcription Factors | 187 | 198 | (Lew, Kapopoulou, Jones, & Cole, 2011)^31^ |
| Trehalose | 10 | 10 | (Wilson et al., 1999)^33^ |
| Triacylglycerol Synthases | 14 | 16 | (Thanna & Sucheck, 2016)^34^ |
| Universal stress proteins | 9 | 10 | (Lew, Kapopoulou, Jones, & Cole, 2011)^31^ |
| Zur regulon | 17 | 20 | (Dow et al., 2021)^35^ |

### 2. **Supplemental Results**

#### **Figure S2.** Flow diagram of design/testing of SEARCH-TB

A custom assay was created using Illumina AmpliSeq technology with the H37RV *Mtb* strain used as a reference a long with eight other *Mtb* reference strains. To avoid off-target amplification of non-*Mtb­* organisms, genes with similar sequences in 12 exclusion genomes were removed. After designing the assay, primers were compared to the Erdman strain and genes which corresponded to primers not effective on the Erdman strain were removed from further analysis. Finally, genes which failed to adequately amplify in quality assurance experiments with genomic DNA were also excluded from downstream analysis. This resulted in 3,568 genes used in the final analysis.

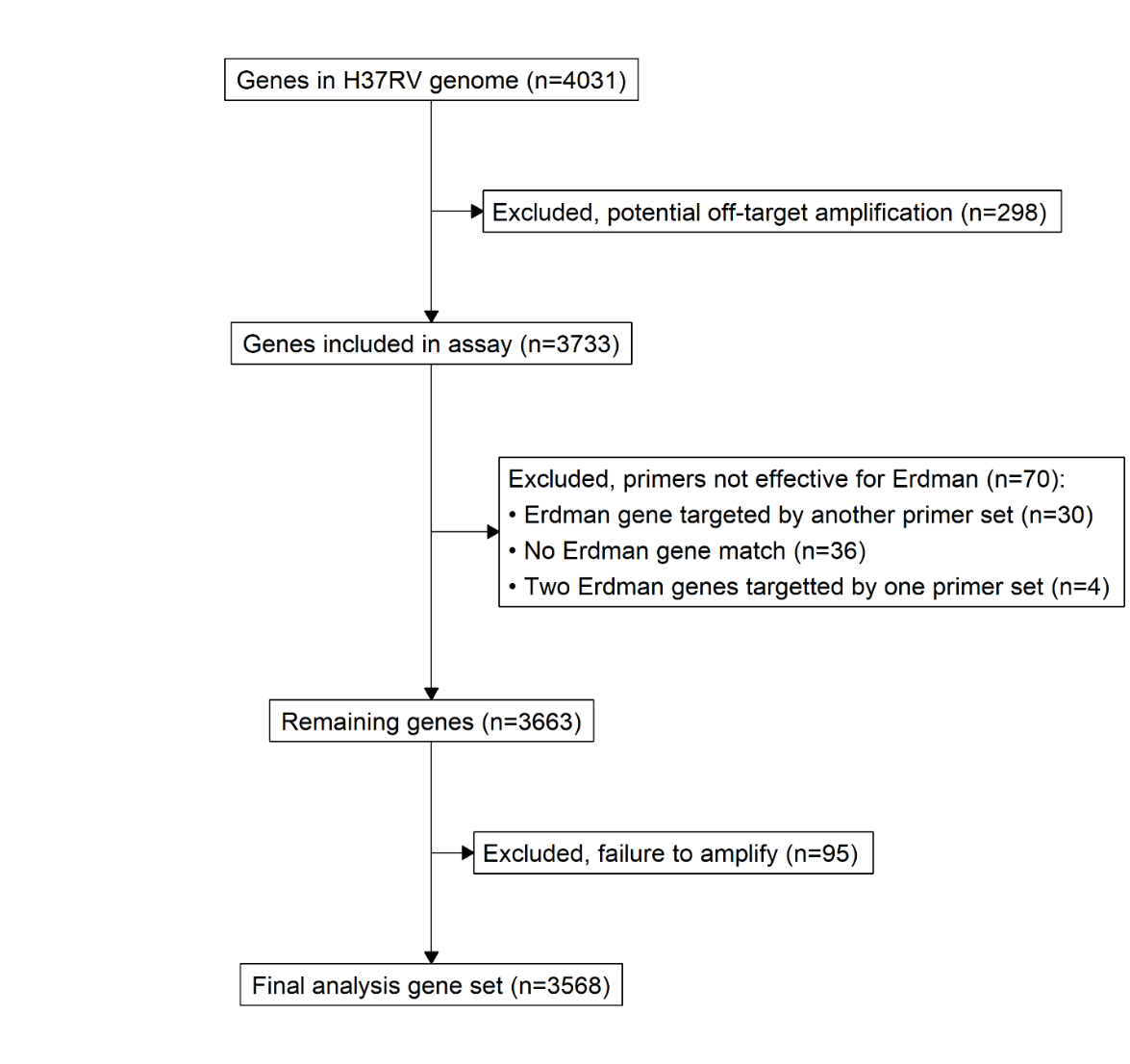

#### **Figure S3.** Comparison batch effect and treatment effect.

As described in the manuscript Methods section, 19 samples underwent library prep and sequencing via SEARCH-TB twice. Batch effect values were calculated by averaging the absolute log_2_ fold difference in gene expression values between replicate pairs across all pairs of replicates. Treatment effect values were extracted from edgeR models comparing treatment groups. This demonstrates that the magnitude of the batch effect was very small relative to the treatment effect.

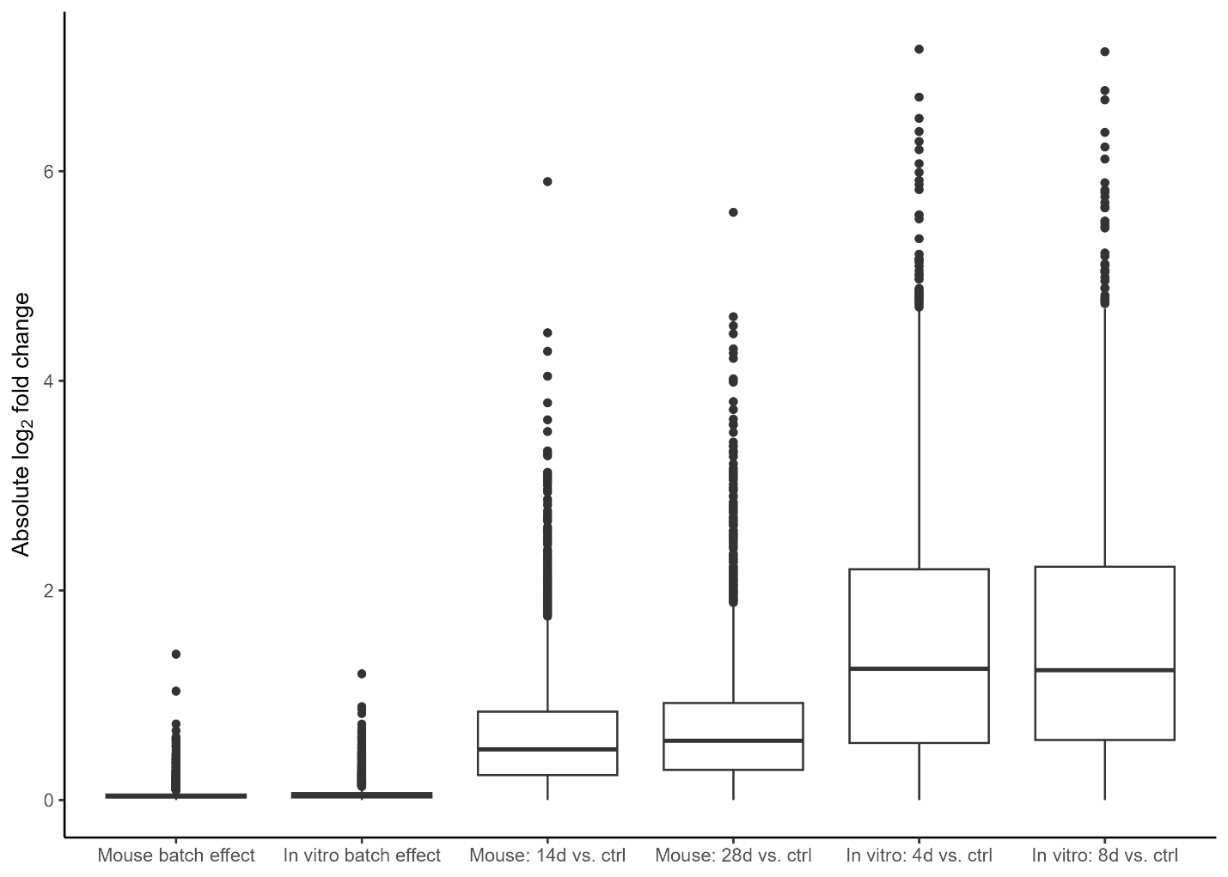

#### **Table S3***.* Expression of efflux pumps.

Fold-change and adjusted *P-*values for efflux pumps with potential drug-transporting capacity as identified in Remm, *et.al.*^17^

| **Transporter Superfamily** | **Pump/Gene RV (symbol)** | **Mouse day 28** | | **In vitro day 8** | |
| --- | --- | --- | --- | --- | --- |
|  |  | **log_2_ FC** | **adj-*P* val** | **log_2_ FC** | **adj-*P* val** |
| RND transporter | MmpL5-MmpS5 |  |  |  |  |
|  | Rv0676c (mmpL5) | -0.34 | **2.79E-02** | -0.12 | 5.59E-01 |
|  | Rv0677c (mmpS5) | -0.67 | **2.59E-03** | 0.66 | **1.79E-02** |
| MFS transporter | Rv1258c (Tap) |  |  |  |  |
|  | Rv1258c | 1.04 | **3.05E-08** | 1.62 | **1.05E-11** |
| SMR transporters | Rv3065 (Mmr) |  |  |  |  |
|  | Rv3065 (mmr) | 0.43 | **4.66E-02** | 0.19 | 4.94E-01 |
| ABC transporter | Rv2686c-2688c |  |  |  |  |
|  | Rv2686c | 1.20 | **4.07E-08** | 2.70 | **4.21E-21** |
|  | Rv2687c | 1.05 | **8.24E-08** | 2.52 | **6.90E-23** |
|  | Rv2688c | 1.22 | **1.27E-11** | 3.31 | **7.67E-41** |
| ABC transporter | Rv0194 |  |  |  |  |
|  | Rv0194 | 0.70 | **4.06E-06** | 2.37 | **9.58E-33** |
| ABC transporter | Rv2936-2938 (DrrABC) |  |  |  |  |
|  | Rv2936 (drrA) | -1.82 | **4.44E-18** | -1.41 | **9.77E-08** |
|  | Rv2937 (drrB) | -1.00 | **1.09E-08** | -0.64 | **4.12E-03** |
|  | Rv2938 (drrC) | -0.45 | **4.41E-03** | 0.70 | **3.55E-04** |
| MFS transporter | Rv1634 |  |  |  |  |
|  | Rv1634 | 0.40 | **1.40E-02** | 1.29 | **2.31E-10** |
| MFS transporter | Rv2333c (Stp) |  |  |  |  |
|  | Rv2333c (stp) |  |  |  |  |
| MFS transporter | Rv0849 |  |  |  |  |
|  | Rv0849 | 0.34 | 9.34E-02 | 1.66 | **5.46E-11** |
| MFS transporter | Rv2846c (EfpA) |  |  |  |  |
|  | Rv2846c (efpA) | -0.66 | **1.29E-04** | -1.12 | **1.49E-07** |
| ABC transporter | Rv1217c-Rv1218c |  |  |  |  |
|  | Rv1217c | 0.78 | **4.75E-05** | 0.94 | **9.75E-05** |
|  | Rv1218c | 0.89 | **2.18E-07** | 0.37 | 9.49E-02 |
| ABC transporter | Rv1456c-Rv1457c-Rv1458c |  |  |  |  |
|  | Rv1456c | 0.09 | 7.61E-01 | -0.59 | 8.48E-02 |
|  | Rv1457c | 0.24 | 2.75E-01 | -0.31 | 2.59E-01 |
|  | Rv1458c | -0.20 | 1.92E-01 | -0.25 | 2.07E-01 |
| PDIM synthesis | Rv2942 (MmpL7) |  |  |  |  |
|  | Rv2942 (mmpL7) | -0.23 | 1.29E-01 | -0.56 | **2.53E-03** |
| MFS transporter | Rv1410c (P55) |  |  |  |  |
|  | Rv1410c | -2.03 | **8.41E-21** | -2.83 | **1.48E-23** |
| ABC transporter | Rv1819c (BacA) |  |  |  |  |
|  | Rv1819c (bacA) | 0.88 | **2.07E-06** | -0.45 | 5.63E-02 |
| ABC-F subfamily | Rv2477c |  |  |  |  |
|  | Rv2477c | -1.90 | **4.23E-16** | -2.36 | **3.59E-15** |
| ABC transporter | Rv0342 and Rv0933 |  |  |  |  |
|  | Rv0342 (iniA) | 0.46 | **1.48E-03** | 1.95 | **8.68E-27** |
|  | Rv0933 (pstB) | -0.34 | 5.14E-02 | -1.44 | **1.82E-11** |

**Table S4***.* Expression of drug targets.

Fold-change and adjusted *P-*values for genes for targets identified by the Working Group on New TB Drugs^15^ and Shetye, *et., al.*^16^

| **Drug or class** | **Gene** | **Symbol** | **log_2_ FC** | **adj-*P-*val** | **log_2_ FC** | **adj-*P-*val** |
| --- | --- | --- | --- | --- | --- | --- |
| **Approved drugs** | | | | | | |
| **Isoniazid** | Rv1908c | katG | -1.9 | **1.6E-07** | -2.8 | **1.3E-09** |
|  | Rv1484 | inhA | -1.1 | **5.9E-09** | -2.3 | **1.1E-19** |
| **Rifamycins** | Rv0667 | rpoB | -0.1 | 7.0E-01 | -1.7 | **2.6E-05** |
| **Pyrazinamide** | Rv2043c | pncA | -0.8 | **1.4E-05** | -0.7 | **3.8E-03** |
| **Ethambutol** | Rv3795 | embB | 0.3 | 9.9E-02 | -0.6 | **8.6E-03** |
| **Fluoroquinolones** | Rv0005 | gyrB | -0.9 | **1.8E-05** | -1.4 | **3.4E-08** |
|  | Rv0006 | gyrA | -1.1 | **2.9E-09** | -2.2 | **7.5E-21** |
| **Diarylquinolones** | Rv1305 | atpE | -3.3 | **7.5E-37** | -4.6 | **1.6E-38** |
| **D-cycloserine** | Rv2981c | ddlA | -0.1 | 4.3E-01 | -0.3 | 1.1E-01 |
| **Ethionamide** | Rv3854c | ethA | -0.3 | 1.1E-01 | 0.8 | **2.5E-05** |
| **Beta-lactams (D, D-transpeptidases)** | Rv2911 | dacB2 | 0.3 | 2.5E-01 | -1.7 | **1.1E-07** |
| **Carbapenems (L, D-transpeptidases)** | Rv0116c | ldtA | -1.0 | **1.4E-07** | -0.1 | 8.4E-01 |
|  | Rv2518c | ldtB | -1.7 | **6.6E-11** | -1.7 | **6.3E-07** |
| **Clofazamine** | Rv1854c | ndh | -1.9 | **2.1E-11** | -1.6 | **3.7E-06** |
| **Investigational** | | | | | | |
| **Peptidoglycan Layer** |  |  |  |  |  |  |
| **Mur ligase inhibitors** | Rv0482 | murB | 0.7 | **1.4E-04** | -0.1 | 6.5E-01 |
|  | Rv1315 | murA | -0.3 | 9.2E-02 | -1.2 | **2.1E-08** |
|  | Rv2152c | murC | 0.3 | **3.8E-02** | 0.3 | 8.3E-02 |
|  | Rv2155c | murD | 0.2 | 4.3E-01 | 0.3 | 3.4E-01 |
|  | Rv2157c | murF | 0.2 | 3.2E-01 | -0.2 | 3.6E-01 |
| **D-alanine:D-alanine Ligase** | Rv2981c | ddlA | -0.1 | 4.3E-01 | -0.3 | 1.1E-01 |
| **Translocase 1 inhibitor** | Rv2156c | murX | 0.2 | 2.2E-01 | -0.6 | **9.4E-03** |
| **MurX inhibitors** | Rv2156c | murX | 0.2 | 2.2E-01 | -0.6 | **9.4E-03** |
| **GlcN-1-p analogues** | Rv1018c | glmU | -0.3 | 4.5E-01 | -0.1 | 7.6E-01 |
| **Polyketide synthase Pks13** | Rv3800c | pks13 | -1.8 | **1.9E-13** | -1.5 | **3.6E-07** |
| **Arabinogalactan Layer** |  |  |  |  |  |  |
| **DprE1 inhibitors** | Rv3790 | dprE1 | 0.1 | 6.9E-01 | 0.2 | 5.2E-01 |
| **Arabinosyltransferase C** | Rv3793 | embC | 0.0 | 8.8E-01 | -0.7 | **5.3E-05** |
| **WecA inhibitor** | Rv1302 | rfe | 0.5 | **2.6E-04** | 1.2 | **2.0E-11** |
| **Mycolic Acid Layer** |  |  |  |  |  |  |
| **MmpL3 inhibitors** | Rv0206c | mmpL3 | -1.1 | **2.4E-08** | -2.4 | **9.3E-21** |
| **KasA inhibitor** | Rv2245 | kasA | -2.7 | **1.7E-18** | -3.7 | **9.7E-20** |
| **Translation** |  |  |  |  |  |  |
| **Leucyl-tRNA synthase inhibitor** | Rv0041 | leuS | 0.1 | 6.7E-01 | -0.4 | **1.9E-02** |
| **DNA replication** |  |  |  |  |  |  |
| **DNA gyrase A inhibitor** | Rv0006 | gyrA | -1.1 | **2.9E-09** | -2.2 | **7.5E-21** |
| **DNA gyrase B inhibitor** | Rv0005 | gyrB | -0.9 | **1.8E-05** | -1.4 | **3.4E-08** |
| **Energy metabolism** |  |  |  |  |  |  |
| **Qcrb inhibitor (Cytochrome bc1-aa3)** | Rv2196 | qcrB | -2.2 | **4.6E-19** | -3.3 | **1.4E-24** |
| **Proteolysis & Proteostasis** |  |  |  |  |  |  |
| **clpC1 inhibitor** | Rv3596c | clpC1 | -1.5 | **2.1E-09** | -3.4 | **2.2E-23** |
| **clpP1 and clpP2 inhibitor** | Rv2460c | clpP2 | -1.7 | **2.2E-13** | -3.6 | **3.2E-31** |
|  | Rv2461c | clpP1 | -1.8 | **1.4E-12** | -3.0 | **2.7E-19** |
| **Cellular metabolism** |  |  |  |  |  |  |
| **Tryptophan synthase inhibitor** | Rv1612 | trpB | -0.3 | 5.7E-02 | -1.8 | **2.9E-16** |
|  | Rv1613 | trpA | -1.1 | **1.8E-09** | -2.0 | **1.3E-17** |
| **Aspartate decarboxylase (PZA)** | Rv3601c | panD | -1.1 | **1.8E-05** | -0.9 | **4.0E-03** |
| **Efflux transport of antibiotics** | Rv2846c | efpA | -0.7 | **1.3E-04** | -1.1 | **1.5E-07** |

### 3. **References**

6.	Andrews, S. FastQC: A Quality Control Tool for High Throughput Sequence Data. (2010).
